## Supplemental Materials Inventory for "Actin dynamics switches two distinct modes of endosomal fusion in yolk sac visceral endoderm cells"

1. **Supplemental Videos**

**Video S1.** Homotypic fusion of late endosomes in yolk sac VE cells

**Video S2.** Heterotypic fusion of late endosomes in yolk sac VE cells

**Video S3.** Homotypic fusion between late endosomes in VE cells treated with 0.1 μM cytochalasin D

**Video S4.** Bridge fusion between late endosomes in VE cells treated with 1 μM cytochalasin D

**Video S5.** Homotypic fusion between late endosomes in VE cells treated with 1 μM jasplakinolide

**Video S6.** Heterotypic fusion between late endosomes in VE cells treated with 0.1 μM cytochalasin D

**Video S7.** Heterotypic fusion between late endosomes in VE cells treated with 1 μM jasplakinolide

2. **Appendix 1**

Free energy dependence on the size and shape of the membrane

Description of shape relaxation

Free energy landscape for small membranes

Free energy landscape for large membranes

Monte-Carlo simulation of the descending $F^{\mathrm{large}}$ landscape

Video S1. **Homotypic fusion of late endosomes in yolk sac VE cells**

Time-lapse recording of homotypic fusion of late endosomes. At 5 min after labeling with Alexa Fluor 488 transferrin, homotypic fusion of late endosomes was observed frequently in the VE cells. The scale bar indicates 10 μm.

Video S2. **Heterotypic fusion of late endosomes in yolk sac VE cells**

Time-lapse recording of heterotypic fusion between late endosomes and lysosomes. At 15 min after labeling with Alexa Fluor488 transferrin, late endosomes often shrank and disappeared from the focal plane as a result of heterotypic fusion of late endosomes with pre-existing lysosomes that were out of the focal plane. The scale bar indicates 10 μm.

Video S3. **Homotypic fusion between late endosomes in VE cells treated with 0.1 μM cytochalasin D**

Time-lapse recording of homotypic fusion of late endosomes. VE cells pretreted with 0.1 μM cytochalasin D for 5 min were labeled with Alexa Fluor 488-transferrin. Homotypic fusion of late endosomes was observed. The scale bar indicates 10 μm.

Video S4. **Bridge fusion between late endosomes in VE cells treated with 1 μM cytochalasin D**

Time-lapse recording of homotypic fusion of late endosomes. VE cells pretreated with 1 μM cytochalasin D for 5 min were labeled with Alexa Fluor 488-transferrin. Homotypic fusion of late endosomes was observed. The scale bar indicates 3 μm.

Video S5. **Homotypic fusion between late endosomes in VE cells treated with 1 μM jasplakinolide**

Time-lapse recording of homotypic fusion of late endosomes. VE cells pretreated with 1 μM jasplakinolide for 5 min were labeled with Alexa Fluor 488-transferrin. Homotypic fusion of late endosomes was observed. The scale bar indicates 10 μm.

Video S6. **Heterotypic fusion between late endosomes in VE cells treated with 0.1 μM cytochalasin D**

Time-lapse recording of heterotypic fusion between late endosomes and lysosomes in VE cells treated with 0.1 μM cytochalasin D. At 15 min after labeling with Alexa Fluor 488-transferrin, VE cells of embryos were observed. The scale bar indicates 10 μm.

Video S7. **Heterotypic fusion between late endosomes in VE cells treated with 1 μM jasplakinolide**

Time-lapse recording of heterotypic fusion between late endosomes and lysosomes in VE cells treated with 1 μM jasplakinolide. At 15 min after labeling with Alexa Fluor 488-transferrin, VE cells of embryos were observed. The scale bar indicates 10 μm.

**Appendix 1**

**Free energy dependence on the size and shape of membrane**

In general, the free energy governing the shape relaxation process of a vesicle (closed lipid membrane) consists of 2 terms: the bending energy and the osmotic energy,

$$F=b-\Delta p\cdot V$$

The bending energy, $b$, is given by

$$b=\frac{\kappa}{2}\int_{surface} c^{2}\cdot da,$$

where $c$ is the mean curvature defined at each point of the surface of the vesicle and the integration is performed all over the vesicle. $\kappa$ is the bending modulus (Helfrich, 1973). The mean curvature has the dimension of the inverse of the length. Suppose a similarity transformation of a membrane with the length scaling ratio $\lambda$, the mean curvature of the transformed membrane is rescaled by $1/\lambda$ and the surface area is rescaled by $\lambda^{2}$. Thus, the bending energy is invariant under a scale change in size.

The osmotic energy is the product of the osmotic pressure difference between the inside and the outside of the membrane, $\Delta p$, and the volume enclosed by the membrane, $V$. The volume depends on both the size and the shape of the membrane. Here we represent the size of a vesicle with its area $A$, since it is invariant under shape changes, and we introduce a rescaled volume $v$ ($v=0\sim1/{6\sqrt{\pi}}$) with

$$V=A\sqrt{A}\cdot v.$$

Thus,

$$F=b-\Delta p\cdot A\sqrt{A}\cdot v.$$

In the above form, a similarity transformation only changes the scale factor term $A\sqrt{A}$. $b$ and$v$ depend only on the shape and are invariant under a scale change in size.

**Description of shape relaxation**

In this study, we consider a process in which 2 spherical vesicles with different areas connect and the integrated closed membrane proceeds shape relaxation. At the onset of the connection, the membrane shape has rotational symmetry for the axis through the centers of the mass of 2 vesicles and the connection point. We assume that the succeeding shape relaxation also holds the rotational symmetry, because some sort of asymmetric force is necessary to violate the rotational symmetry, which is a more complicated problem than can be discussed here.

For further simplification, we only consider membrane shapes that have a constricted *neck* region, and introduce 2 shape parameters: the neck width, $w$, and the area ratio between both sides of the neck, $a$. The constricted neck region is the trace of the connection event and the time evolution of the 2 parameters expresses the approximate deformation processes of 2 vesicle fusions as follows. Just after fusion, $w=0$ and $a$ is the area ratio between the 2 vesicles. The relaxation of the neck (“explosive” fusion) is expressed by the increase in $w$. The absorption of smaller vesicle by the larger one (“bridge” fusion) is expressed by the decrease in $a$, while $w$ remains small.

Membrane shapes confined by the shape parameters still display a variety. We specify the typical shape of the membrane for a given set of shape parameters $\left( w,a \right)$. The typical shape must be more energetically stable than atypical shapes having the same shape parameter values. We choose the shape having the minimum free energy as the typical shape. Then, we obtain the free energy of the typical shape for each set of shape parameters and display the free-energy landscape on the two-dimensional shape parameter space.

**Free energy landscape for small membranes**

Suppose a considerably small membrane, which is described by small $A\sqrt{A}$, the free energy is approximated by

$$F\simeq F^{\mathrm{small}}=b.$$

To generate rotationally symmetric shapes and to evaluate the bending energy, we consider a curve in the radial and axial space and parametrize it by the function of angle, $\theta$, against the arc-length along the curve, $s$ (Appendix 1-figure 1A). Rotating the curve around the axis gives a membrane shape. The mean curvature is $\frac{d\theta}{ds}+\frac{\sin\left( \theta\right)}{r}$ (Seifert et al., 1991) and the bending energy for the shape is $b=2\pi\kappa\int_{0}^{s_{1}} \frac{r}{2}\left\{ \frac{d\theta}{ds}+\frac{\sin\left( \theta\right)}{r} \right\}^{2}ds.$

We independently determine the shapes giving the minimum bending energy for the right and left parts of the membrane with an additional boundary condition: a smooth connection between the angles at the boundary ($\theta=\frac{\pi}{2}$ at the boundary).

Here we suppose a membrane with the radius of opening $\bar{w}$, area $1$, and give the free energy

$$F= \pi\int_{0}^{s_{1}} L\left( \theta,\theta^{'},r,r^{'},\gamma\right)ds$$

With the Lagrange function

$$L=\kappa r\left\{ \theta^{'}+\frac{\sin\left( \theta\right)}{r} \right\}^{2}+\gamma\left( r^{'}-\cos\left( \theta\right) \right),$$

where $\gamma$ is a Lagrange multiplier. Then the Euler-Lagrange equations for the system are

$$r\frac{d^{2}\theta}{ds^{2}}+\cos\left( \theta\right)\frac{d\theta}{ds}-\frac{\sin\left( \theta\right)\cos\left( \theta\right)}{r}-\gamma\sin\left( \theta\right)=0; \frac{d\gamma}{ds}=\frac{1}{2}\left( \frac{d\theta}{ds} \right)^{2}-\frac{\sin^{2} \left( \theta\right)}{2r^{2}}; \frac{dr}{ds}=\cos\left( \theta\right).$$

Solving these equations with boundary conditions $\left( \theta,r \right)_{s=0}=\left( 0,0 \right)$, $\left( \theta,r \right)_{s=s_{1}}=\left( \frac{\pi}{2},\bar{w} \right)$ and $2\pi\int_{0}^{s_{1}} rds=1$, we get the rotationally symmetric shape having a minimum bending energy $\theta=\theta_{\min}\left( s;\bar{w} \right)$.

We calculated the minimum energy shape for $\bar{w}\in\left[ 0,1/\sqrt{2\pi} \right]$, and plotted the bending energy against the radius of the opening. The function is well fitted with a hyperbolic curve

$$\bar{b}\left( \bar{w} \right)=4\pi\kappa\left\{ \frac{\sqrt{\lambda^{2}\left( \sqrt{2\pi}\bar{w}-1 \right)^{2}+1}-1}{\sqrt{\lambda^{2}+1}-1}+1 \right\}$$

with $\lambda\simeq7.467$ (Appendix 1-figure 1C). Thus, we generate the minimum-bending-energy shape for a given set of shape parameters by combining two rescaled partials spheres with scaling ratios $a$ and $1-a$, and we get the free energy as a function of $\left( w,a \right)$

$$F^{\mathrm{small}}\left( w,a \right)=b\left( w,a \right)=\bar{b}\left( \frac{w}{\sqrt{a}} \right)+\bar{b}\left( \frac{w}{\sqrt{1-a}} \right)$$

**Free energy landscape for large membranes**

Contrary to the small membrane, a considerably large membrane is described by a large $\Delta P$. We tentatively approximate the free energy of the large membrane with

$$F\simeq F_{0}^{\mathrm{large}}=-\Delta p\cdot A\sqrt{A}\cdot v.$$

The equation indicates that the minimum-free-energy shape corresponds to the shape with the largest volume for the given shape parameters $\left( w,a \right)$. Moreover, we independently determine the shapes giving the maximum volume for the right and left parts of the membrane across the constricted neck region, since the area ratio and the size of the boundary (the constricted neck region) are determined by shape parameters. The shape of each part of the membrane corresponds to a partial sphere (Appendix 1-figure 1B).

A general partial sphere with the radius of opening $\tilde{w}$, area $1$, and the volume $\tilde{v}$, is given by

$$\tilde{w}=\alpha\sin\left( \psi\right)$$

$$1=2\pi\alpha^{2}\left( 1-\cos\left( \psi\right) \right)$$

$$\tilde{v}=\pi\alpha^{3}\left\{ \frac{\cos^{3} \left( \psi\right)}{3}-\cos\left( \psi\right)+\frac{2}{3} \right\}$$

where, $\alpha$ and $\psi$ are the radius and angle of the partial sphere, respectively (Appendix 1-figure 1B). Solving the volume as a function of $\tilde{w}$, we get

$$\cos\left( \psi\right)=2\pi\tilde{w}^{2}-1, \alpha=\frac{1}{2\sqrt{\pi\left( 1-\pi\tilde{w}^{2} \right)}}$$

$$\tilde{v}\left( \tilde{w} \right)=\frac{1}{8\pi\left( 1-\pi\tilde{w}^{2} \right)\sqrt{\pi\left( 1-\pi\tilde{w}^{2} \right)}}\left\{ \frac{\left( 2\pi\tilde{w}^{2}-1 \right)^{3}}{3}-2\pi\tilde{w}^{2}+\frac{5}{3} \right\}.$$

We generate the maximum volume shape for a given set of shape parameters by combining two rescaled partials spheres with scaling ratios $\sqrt{a}^{3}$ and $\sqrt{1-a}^{3}$, and the volume of the shape is calculated by

$$v\left( w,a \right)=\sqrt{a}^{3}\cdot\tilde{v}\left( \frac{w}{\sqrt{a}} \right)+\sqrt{1-a}^{3}\cdot\tilde{v}\left( \frac{w}{\sqrt{1-a}} \right)$$

Therefore, we get the free energy as a function of $\left( w,a \right)$

$$F_{0}^{\mathrm{large}}\left( w,a \right)=-\Delta P\cdot\left\{ \sqrt{a}^{3}\cdot\tilde{v}\left( \frac{w}{\sqrt{a}} \right)+\sqrt{1-a}^{3}\cdot\tilde{v}\left( \frac{w}{\sqrt{1-a}} \right) \right\}$$

The resulting shape of the membrane has a sharply bent region at the neck, which stores large bending energy. Moreover, at the onset of fusion, the osmotic energy has no gradient in the $w$ direction

$$\left. \frac{\partial F_{0}^{\mathrm{large}}\left( w,a \right)}{\partial w} \right|_{w=0}=0.$$

Thus, the above approximation of neglecting the bending energy for large membranes is not always valid. To draw the time evolution of $w$, we have to incorporate the bending energy as a modification term.

Suppose that the neck region is curved with radius $R_{0}$ (Appendix 1-figure 1D and E) which is considerably smaller than the actual neck width $W=\sqrt{A}w$, the bending energy of the neck region is

$$B^{neck}=2\pi\kappa\int_{\left( \pi-\psi_{1} \right)r_{0}}^{\psi_{2}r_{0}} \frac{W-R_{0}\sin\left( \xi\right)}{2}\left\{ \frac{\partial\xi}{\partial s}+\frac{\sin\left( \xi\right)}{W-R_{0}\sin\left( \xi\right)} \right\}^{2}ds,$$

with $s=r_{0}\theta, \xi=\pi-\theta, {\partial\xi}/{\partial s}={\partial\xi}/{\partial\theta}\cdot{\partial\theta}/{\partial s}={-1}/{r_{0}}$, and the volume change by introducing the curved neck region is negligible. The coordinate system of the calculation is shown in Appendix 1-figure 1D and E. Using the relation $W\gg R_{0}$, the bending energy becomes

$$B^{neck}\simeq2\pi\kappa\int_{\pi-\psi_{1}}^{\psi_{2}} \frac{W}{2}\left\{ \frac{-1}{R_{0}} \right\}^{2}R_{0}d\theta=2\pi\kappa\frac{W}{R_{0}}\left( \mathrm{acos} \left( \frac{2\pi w^{2}}{a}-1 \right)+\mathrm{acos} \left( \frac{2\pi w^{2}}{1-a}-1 \right)-\pi\right).$$

Since $R_{0}$ is assumed to be the minimum curvature radius determined by the physical properties of the membrane, it should be independent from the scale change in size (in other words, the thickness of the neck is fixed under the scale change). Therefore, the bending energy of the neck region depends on the size. The right-hand side of the above equation is decomposed into the size-dependent parameter and the size-independent function of $\left( w,a \right)$,

$$\Gamma=2\pi\kappa\frac{\sqrt{A}}{R_{0}}, b^{neck}\left( w,a \right)=w\left\{ \mathrm{acos} \left( \frac{w^{2}}{a}-1 \right)+\mathrm{acos} \left( \frac{w^{2}}{1-a}-1 \right)-\pi\right\}.$$

Finally, we get the modified free energy for the large membrane

$$F^{\mathrm{large}}\left( w,a \right)=\Gamma\cdot b^{neck}\left( w,a \right)-\Delta p\cdot A\sqrt{A}\cdot v\left( w,a \right).$$

Since $\Gamma\propto\sqrt{A}$, the contribution of the osmotic energy is dominant for the large membrane except the region where the slope of the osmotic energy vanishes.

**Monte-Carlo simulation descending** $\boldsymbol{F}^{\mathbf{large}}$ **landscape**

Monte-Carlo simulations on $\left( w,a \right)$ space were performed with free energy $F^{\mathrm{large}}\left( w,a \right)$. We assumed that active fluctuations caused by nonequilibrium processes or activities of motor proteins are modeled by thermal fluctuations with shifted temperature (effective temperature) (Ben-Isaac et. al., 2011). Although this system has three parameters—the size of the vesicle $\sqrt{A}$, osmotic pressure difference $\Delta p$, and energy of the effective temperature $\Xi$—it is invariant under the transformation $\left( \sqrt{A},\Delta p,\Xi\right)\to\left( \lambda\sqrt{A},{\Delta p}/{\lambda^{2}},\lambda\Xi\right).$ It means that the essential dimension of the parameters is two. Thus, we chose two rescaled parameters ${\Delta pV}/\Gamma$ and $\Xi/\Gamma$ with $V=4A\sqrt{A}/{3\sqrt{\pi}}$, $\Gamma=4\pi\kappa\sqrt{A}/{R_{0}}$ to be changed. With the initial condition $\left( w,a \right)=(0.0,0.3)$, Monte-Carlo simulations are performed 1000 times for each set of rescaled parameters using the Metropolis method and the frequency of neck expansions in the courses of shape relaxations was calculated. Neck expansion was judged when $w=0.2$.
