## Supplementary figures and images for "Actin dynamics switches two distinct modes of endosomal fusion in yolk sac visceral endoderm cells"

### Figure 1-figure supplement 1

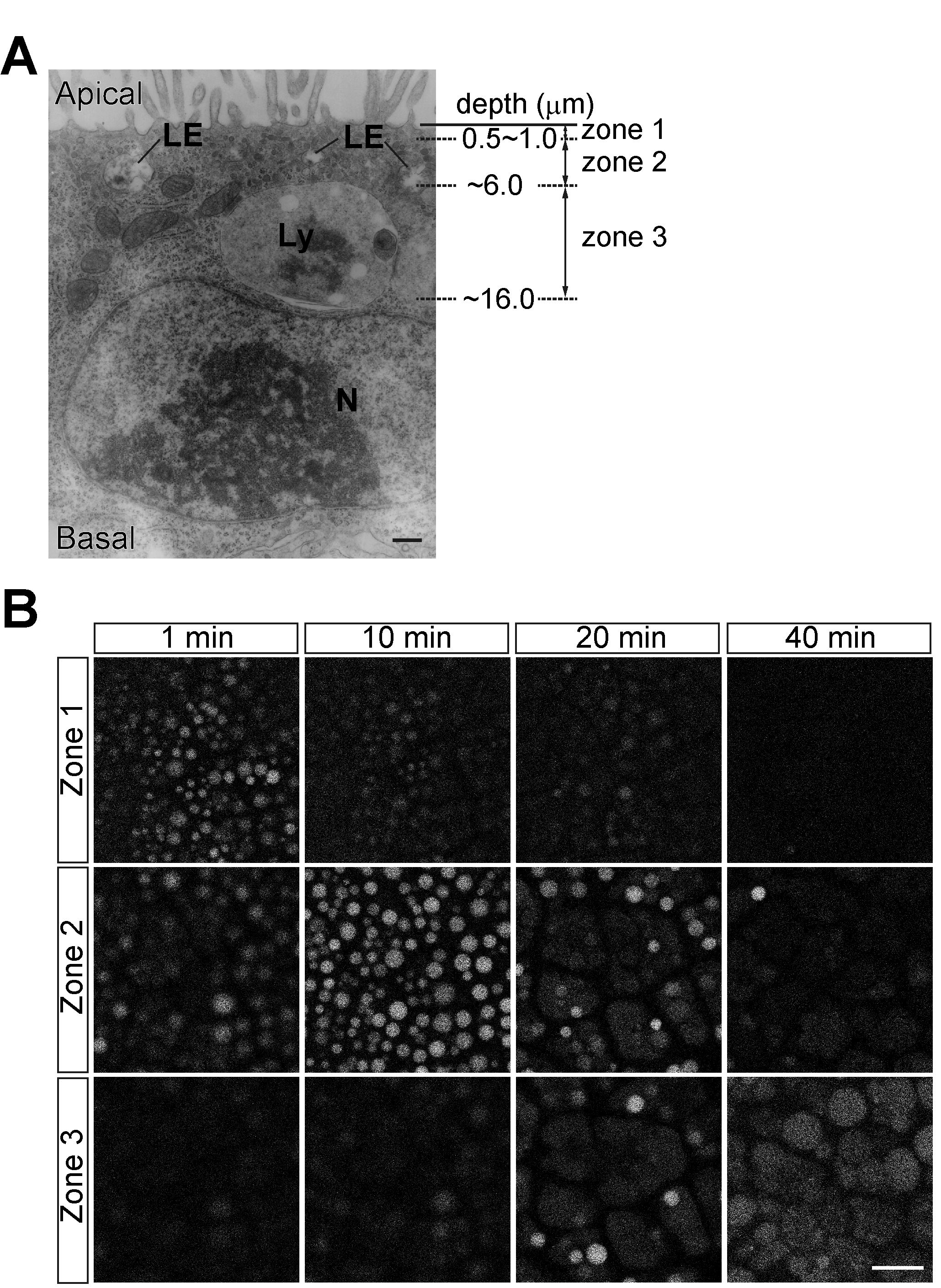

### Figure 4-figure supplement 1

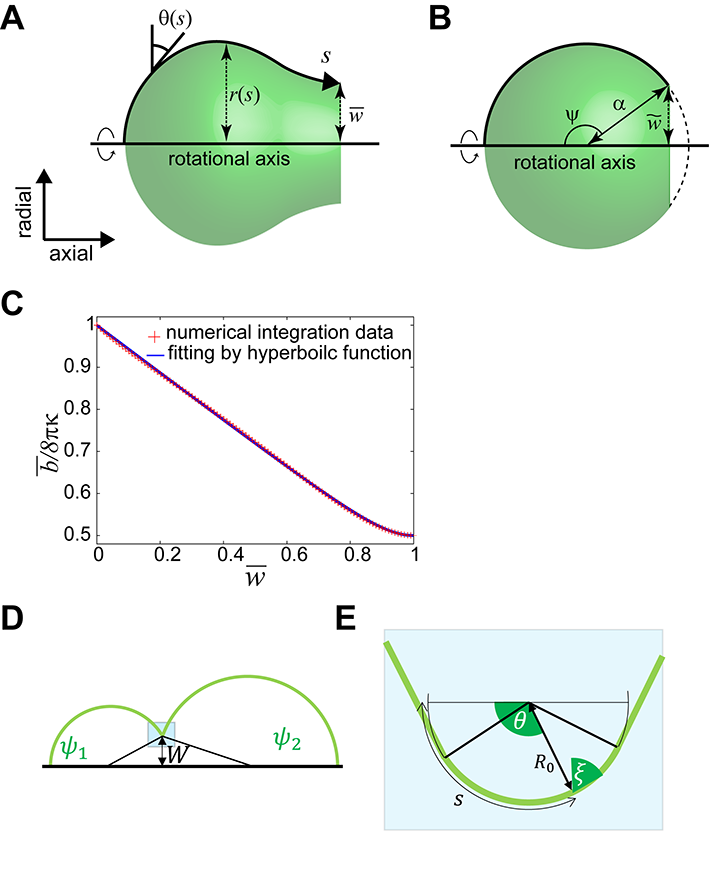

### Figure 6-figure supplement 1

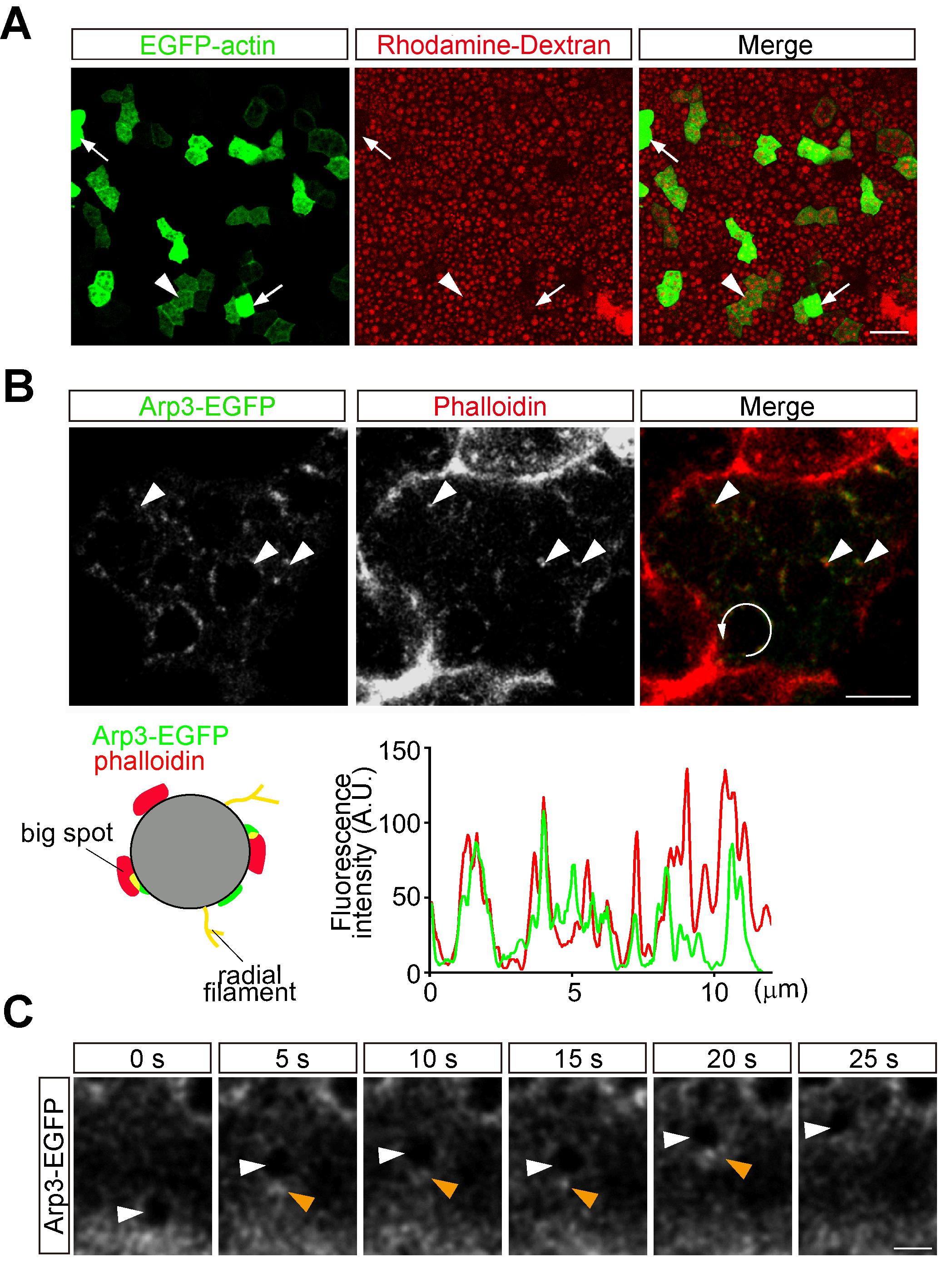

### Figure 7-figure supplement 1

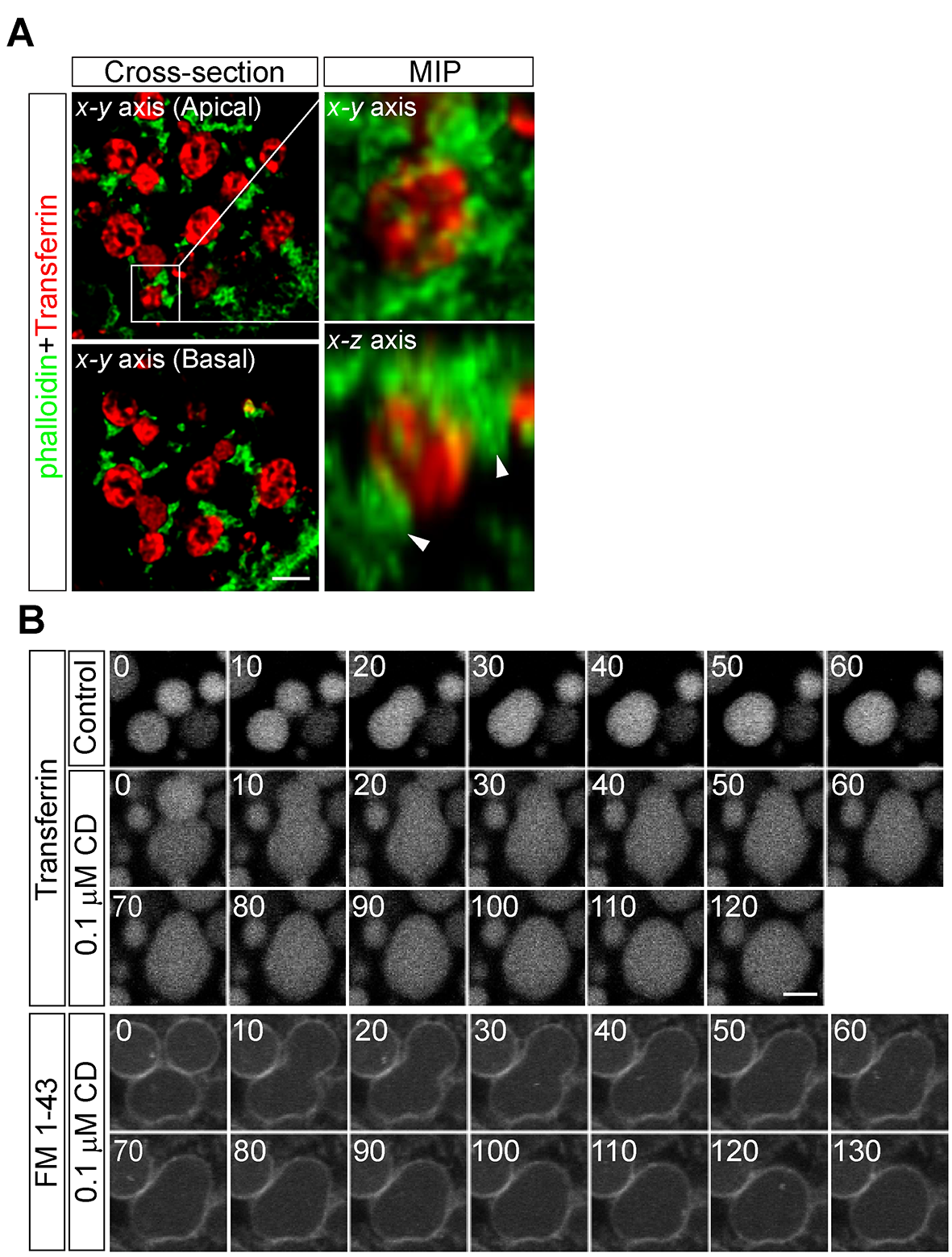

### Figure 7-figure supplement 2

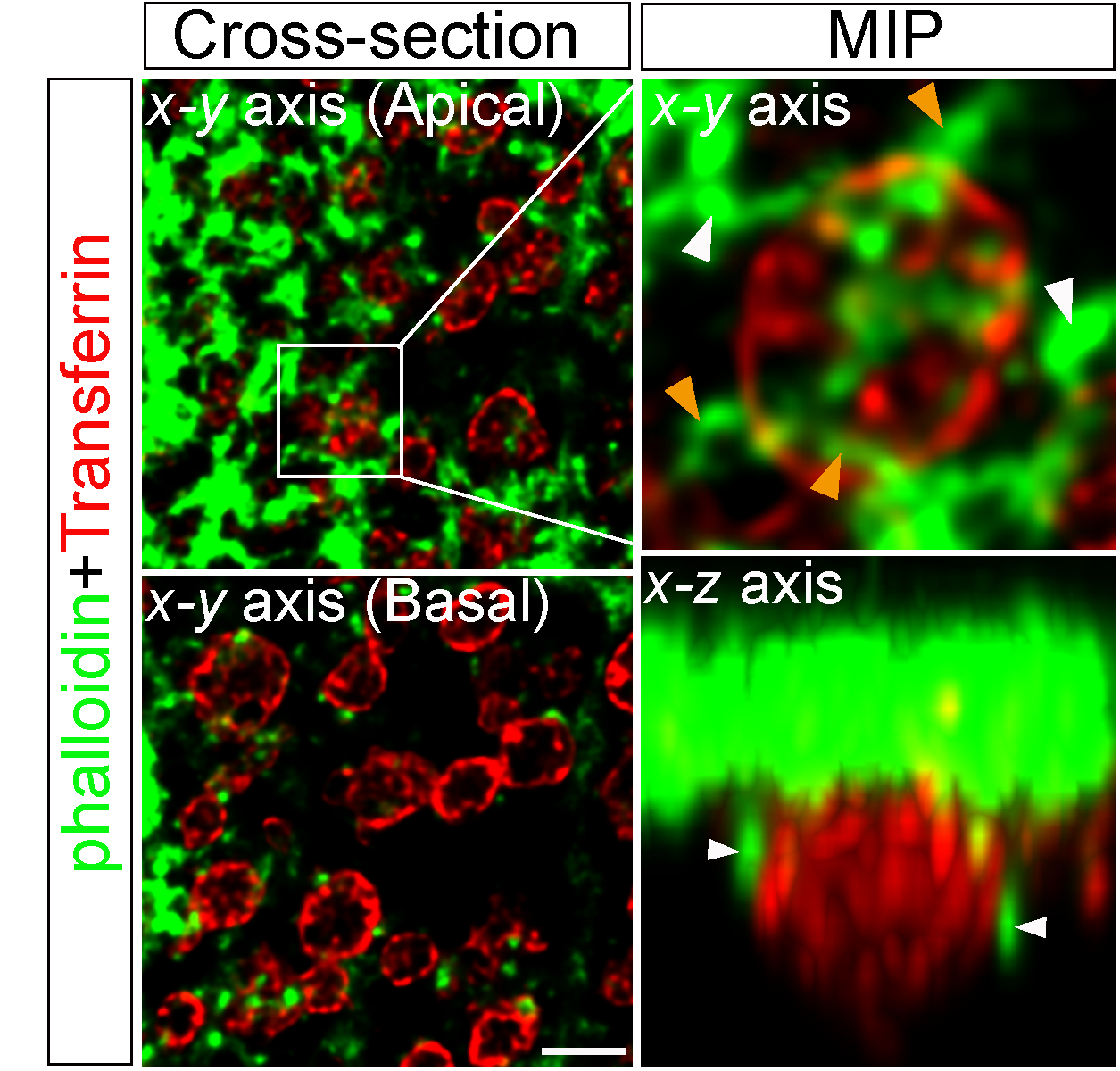

### Figure 7-figure supplement 3

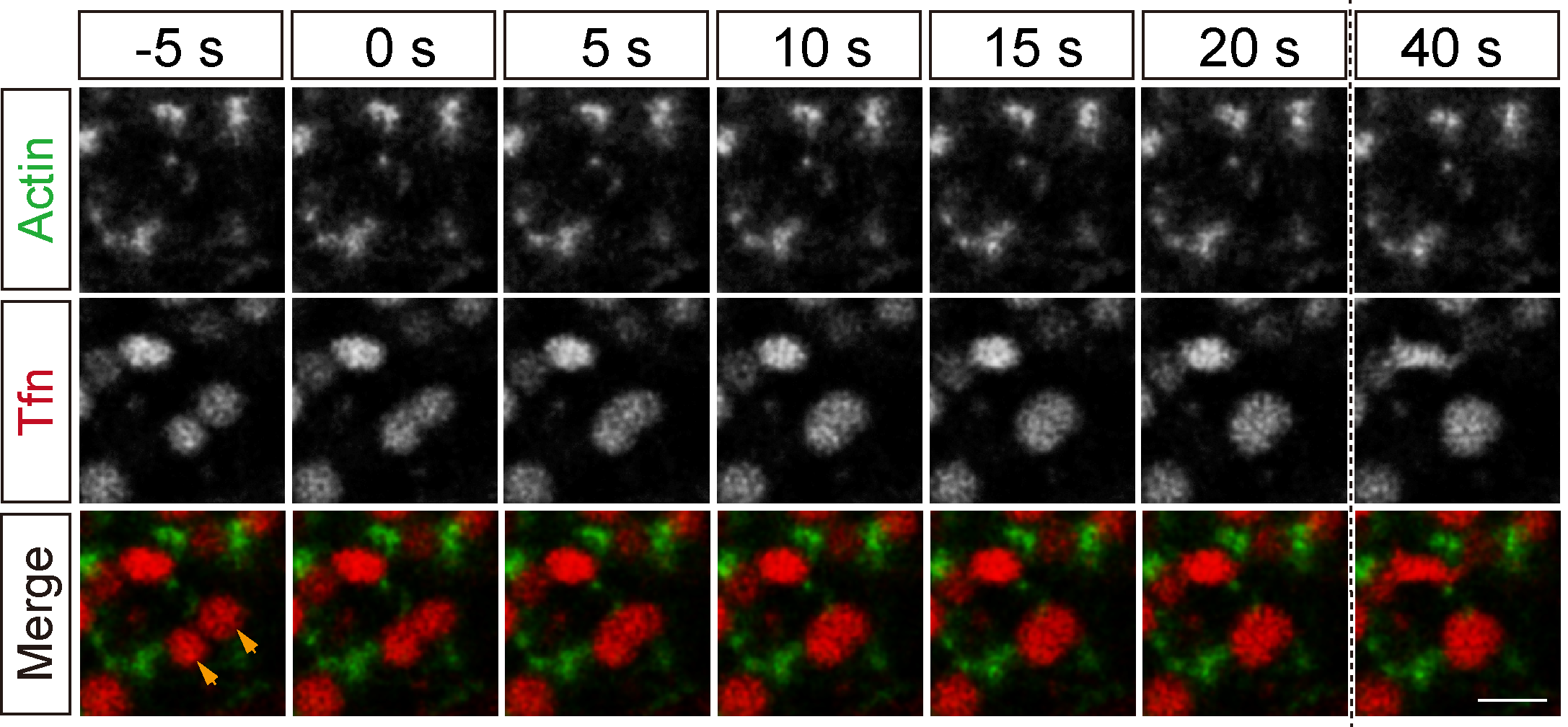

### Figure 7-figure supplement 4

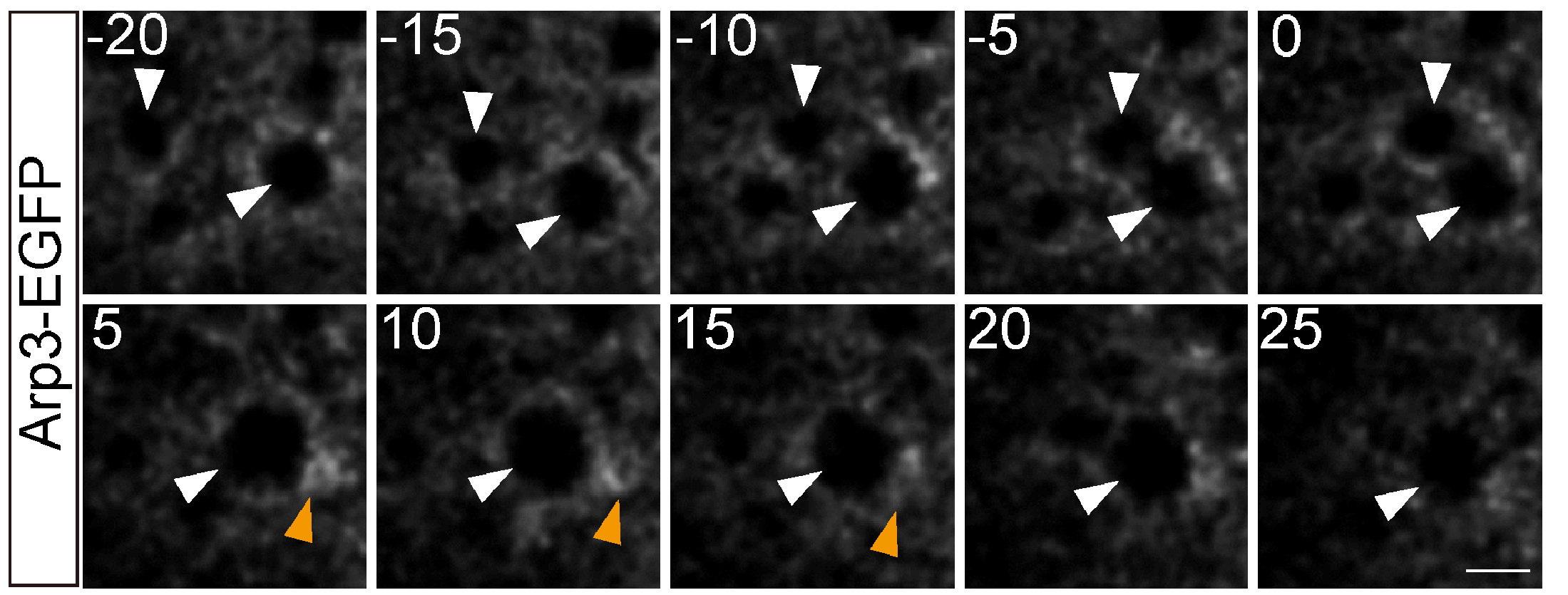

### Figure 8-figure supplement 1

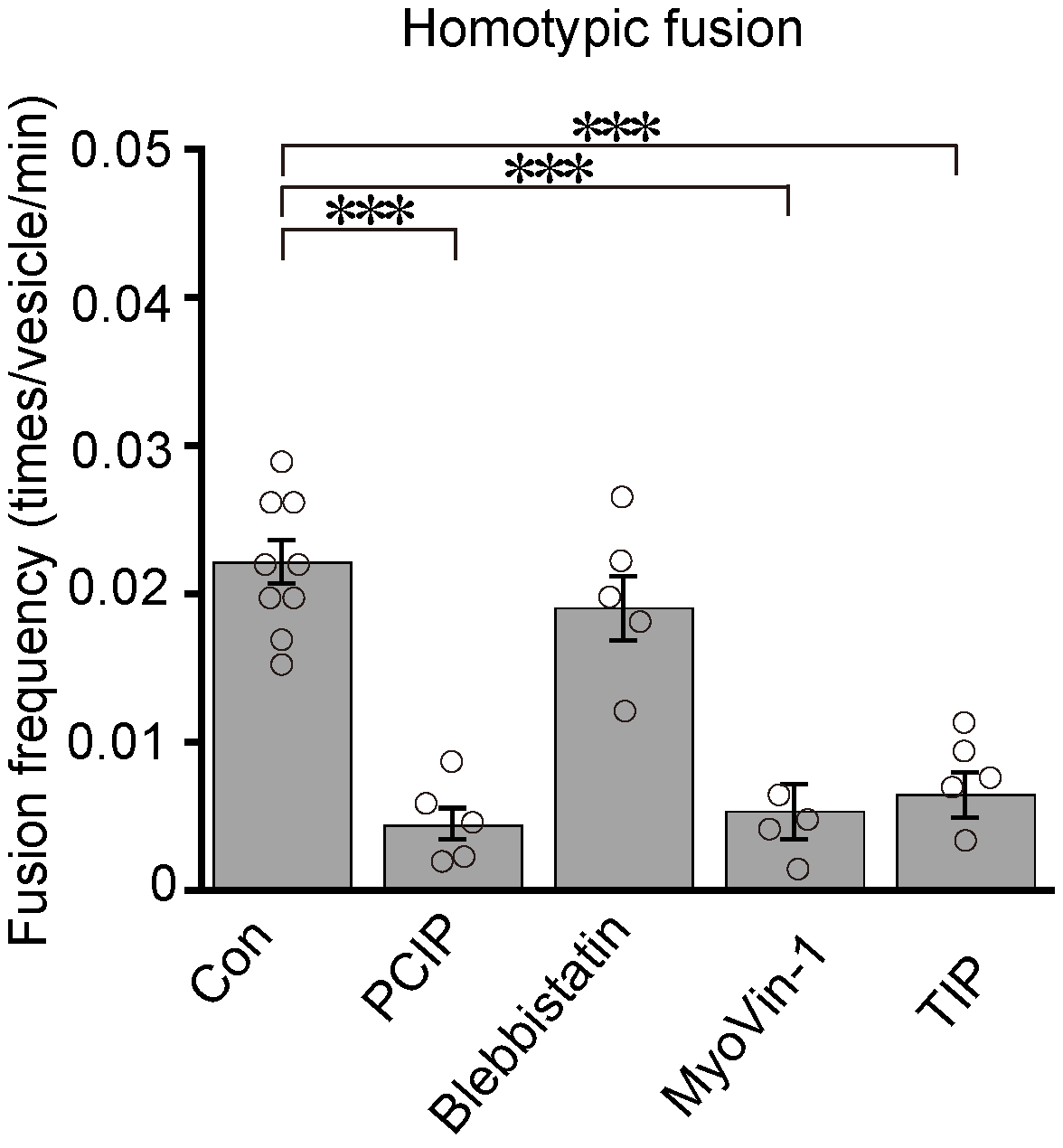

### Figure 9-figure supplement 1

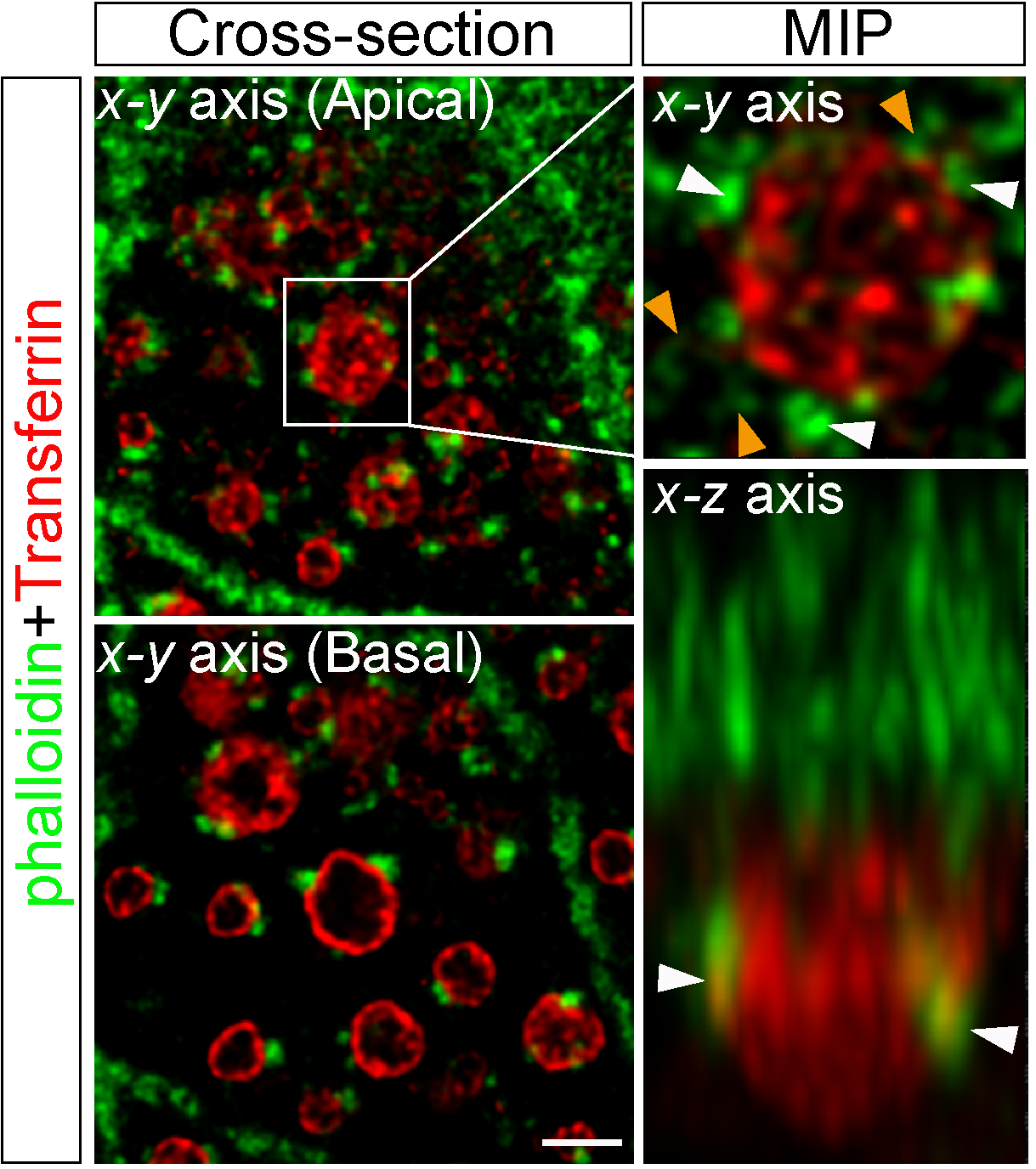
